# Dual RNA-seq reveals large-scale non-conserved genotype x genotype specific genetic reprograming and molecular crosstalk in the mycorrhizal symbiosis

**DOI:** 10.1101/393637

**Authors:** Ivan D. Mateus, Frédéric G. Masclaux, Consolée Aletti, Edward C. Rojas, Romain Savary, Cindy Dupuis, Ian R. Sanders

## Abstract

Arbuscular mycorrhizal fungi (AMF) impact plant growth and are a major driver of plant diversity and productivity. We quantified the contribution of intra-specific genetic variability in cassava (*Manihot esculenta*) and *Rhizophagus irregularis* to gene reprogramming in symbioses using dual RNA-sequencing. A large number of cassava genes exhibited altered transcriptional responses to the fungus but transcription of most of these plant genes (72%) responded in a different direction or magnitude depending on the plant genotype. Two AMF isolates displayed large differences in their transcription, but the direction and magnitude of the transcriptional responses for a large number of these genes was also strongly influenced by the genotype of the plant host. This indicates that unlike the highly conserved plant genes necessary for the symbiosis establishment, plant and fungal gene transcriptional responses are not conserved and are greatly influenced by plant and fungal genetic differences, even at the within-species level. The transcriptional variability detected allowed us to identify an extensive gene network showing the interplay in plant-fungal reprogramming in the symbiosis. Key genes illustrated that the two organisms jointly program their cytoskeleton organisation during growth of the fungus inside roots. Our study reveals that plant and fungal genetic variation plays a strong role in shaping the genetic reprograming in response to symbiosis, indicating considerable genotype x genotype interactions in the mycorrhizal symbiosis. Such variation needs to be considered in order to understand the molecular mechanisms between AMF and their plant hosts in natural communities.

## Introduction

Arbuscular mycorrhizal fungi (AMF) are soil microorganisms that are present in terrestrial ecosystems worldwide [1] and that associate with approximately 74% of land plants [2]. Arbuscular mycorrhizal fungi enhance plant nutrient uptake, plant community diversity and productivity [3], confer plant protection against pathogens and herbivores [4] and protection against abiotic factors such as drought [5] and salt stress [6].

In nature, an individual AMF encounters a variety of different species and genotypes of plants. In natural communities, an individual AMF does not appear to show strong preferences for a given plant species or plant genotype [7]. Despite this, plant species respond in different ways to colonization by the same fungus and the plant response spans the range from positive to negative [8]. Moreover, even different sibling fungi of the same AMF parent, which must be genetically very close to each other, can lead to enormous variation in plant growth [9].

In order to understand the molecular mechanisms of the mycorrhizal symbiosis, the identification of genes involved in the symbiosis has largely relied on forward genetics [10]. This approach relies on identifying the genetic cause of a clear qualitative phenotype. Researchers have mostly focussed on plant mutants in which impaired fungal development occurs. Most of these studies have been conducted at the very early stages of the association, in many cases for less than 21 days post-inoculation, even though the symbiosis lasts for the lifetime of the plant [11]. This approach has uncovered plant genes that are crucial for the colonization and development of the fungus inside the plant; the so-called “mycorrhizal symbiosis toolkit”. These genes are extremely highly conserved throughout much of the plant kingdom [12]. However, using a forward genetics approach with plant mutants that exhibit impaired fungal development, limits the discovery of genes to those that are essential for the establishment of the symbiosis. The fact that the mycorrhizal toolkit of plant genes is highly conserved among plant species that are capable of forming the mycorrhizal symbiosis is highly consistent with experimental and ecological studies indicating that there is little specificity in the symbiosis; in other words, the genetic programing exists in most terrestrial plants to host mycorrhizal fungi. But the approach used to find these genes has also lead to the widespread belief that the plant controls the fungus. There may be a bias towards this assumption based on the particular mutant phenotypes that have been chosen for study, i.e. plant mutants that lead to impaired fungal development.

However, ecological studies show that following establishment of the symbiosis, plant species vary enormously in their growth response to a given AMF taxa [3,8,13] or isolate [14-18]. In addition, different genotypes of the same plant species sometimes, but not always, respond differently to inoculation with a given AMF isolate [19-21]. Thus, the beneficial outcome of the association for the plant is not a foregone conclusion just because upregulation of the symbiotic toolkits of genes occurs and the symbiosis becomes physically established. Thus, while we would expect expression of a conserved repertoire of ‘mycorrhizal toolkit’ genes occurs in all such interactions, we also predict a complex underlying pattern of molecular reprogramming to occur that is plant host genotype and fungal genotype specific.

While significant progress has been made in discovering plant genes that are in involved in the establishment of the symbiosis, identification of important fungal genes involved in these interactions are extremely limited [22-24]. Firstly, this is because stable transformation of AMF has not yet been achieved [25]. Secondly, AMF cannot be cultured without a host. Thirdly, the use of plant mutants that impair fungal development preclude the detection of fungal genes involved in the symbiosis. Finally, because of the lack of knowledge of fungal gene regulation in the symbiosis how plant and fungal genes are co-orchestrated during the symbiosis remains to be elucidated.

Most molecular studies have only evaluated plant transcriptional responses to the fungus by inoculating one plant cultivar with one AMF isolate and comparing the transcriptional response to a mock-inoculated control, without the fungus. Consequently, there is no information on how plant or fungal genetic variation affects plant transcriptional responses to the fungus. Furthermore, there are very few examples where the fungal transcriptional response to different conditions has been measured [26], and to our knowledge, there are no studies that measured the gene transcriptional variation of both partners within the same experimental design. Because of this, it has also not been possible to study how variation in gene transcription of plant genes is related to variation in gene transcription of the AMF partner.

In this study, we aimed to understand the contribution of intra-specific genetic variation of a host plant and an AMF species to the response of each partner at the gene transcription level. In addition, we aimed to identify plant and fungal molecular interactions that are co-orchestrated during the symbiosis. These aims are ecologically relevant because in nature AMF consistently encounter genetically variable partners, and *vice-versa,* and yet the molecular implications of this for the symbiosis are completely unexplored. Understanding how much the molecular interactions between plant and fungus can be shaped by the plant or fungal genotype has been ignored up to now, although the assumption that plant molecular responses to AMF colonization are so strongly conserved has probably hindered advances in our understanding. Consequently, we established a randomized experimental design comprising five genetically different cultivars of *Manihot esculenta*, each inoculated with two different *Rhizophagus irregularis* isolates and a mock-inoculated control. We measured the plant growth response and AMF colonization in each of the treatments. In addition, we performed a dual RNA-seq experiment, to obtain a genome-wide gene transcription profiling of each plant-AMF combination. This combined dataset allowed us to test quantitatively how plant and fungal gene-transcription are affected by the intra-specific variability of the partner organism. In addition, it allowed us to detect plant and fungal genes that are co-transcribed. Such analyses can only be possible with a dataset in which significant variation in both fungal and plant transcripts is the result of different experimental treatments.

We hypothesized that underneath a common “mycorrhizal toolkit” of conserved genes that determines whether the symbiosis will be established, there is considerable non-conserved transcriptional variation of co-regulated genes in the plant and the fungus that is manifested in associations of varying benefit to the plant. Given that the overriding interest in the symbiosis is due to the potential of the fungus to greatly improve plant growth, the lack of knowledge about gene interactions between the two partners represents a fundamental lacuna in our much-needed understanding of the molecular genetics of this symbiosis.

We chose Cassava as the host plant because it is a plant that is known to respond strongly to AMF inoculation in the field [27]. Cassava is considered a vital crop for food security that provides an important source of calories in tropical countries [28]. A good quality cassava genome assembly has recently been published [29]. In addition, cassava can be grown clonally which reduces within-cultivar genetic variation that facilitates the detection of between-cultivar variation. We also focused on *Rhizophagus irregularis* as AMF partner, as it is the main AMF model species with publicly available genome and transcriptome sequences [30-33]. This fungus has been shown to promote the growth of several plants [14], including cassava [34].

## Materials & Methods

### Biological material and growth conditions

We designed a two-factor experiment, where we used five cassava (*Manihot esculenta* Crantz) cultivars inoculated independently with two different mycorrhizal fungi or a mock-inoculated control. The five cassava cultivars were CM6438-14, COL2215, BRA337, CM4574-7 and CM523-7. We carried out a greenhouse experiment with a randomized design to avoid block effects (Supplementary note S1).

### Trait measurements and statistical analyses

After 18 weeks following inoculation, plant height was measured. We then collected and dried the plants for eight days at 72°C. Dry weight of the shoots, bulking roots and fine roots was measured separately. We also randomly selected some fine roots to measure colonization by AMF (Supplementary note S2). We used a binomial generalized linear model for the analysis of the fine root colonization measurements. For all the other quantitative trait variables, we used a mixed model including block as a random effect and tested the significance of the fixed effect of the cassava cultivar, the AMF treatment and their interaction.

### Dual RNA-seq and bioinformatic analysis

We randomly selected several fine root samples from each plant. We extracted RNA, prepared the libraries and sequenced the samples (Supplementary note S3). We then processed the raw reads and separated *in-silico* the cassava and fungal reads (Supplementary figure 1; Supplementary note S4; refer to Supplementary table S1 for the sequenced reads per sample). We then performed a differential expression analysis of each gene (Supplementary note S5) and identified orthologous genes involved in the mycorrhizal symbiosis in *M*. *esculenta* (Supplementary note S6). The genome assembly and annotation available for cassava (Manihot_esculenta_v6.1) was obtained from the plant genomics resource Phytozome (https://phytozome.jgi.doe.gov/). The *R*. *irregularis* gene repertoire was predicted (Supplementary note S7, Supplementary data 1). Finally, we performed gene ontology analysis, functional classification and inferred pathway diagrams of the differentially transcribed genes (Supplementary note S8).

**Figure 1.**
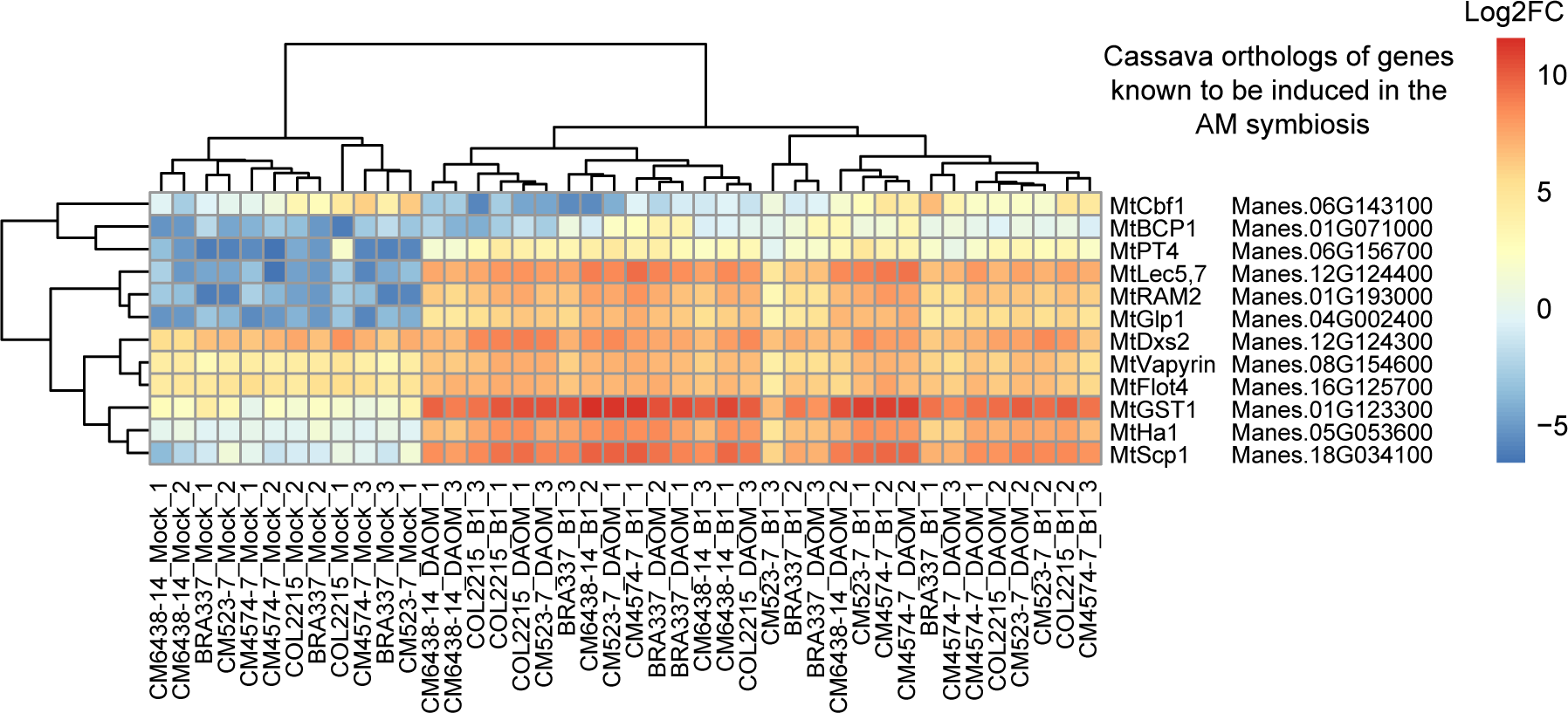
Transcription levels of orthologous genes in inoculated and mock-inoculated cassava that were previously shown to be involved in the establishment of the arbuscular mycorrhizal symbiosis. Genes were chosen from Hogekamp & Küster, 2013 (Supplementary table 4). We clustered the samples by their similarity in log2-counts per million of each gene and by their similarity in transcription across samples.

### Gene co-transcribed networks between cassava and AMF and plant quantitative traits

We used a weighted gene co-expression network analysis (WGCNA) [35] in order to separately build cassava and *R*. *irregularis* gene modules (clusters of genes displaying similar correlated patterns of transcription). As, we were interested in genes involved in the symbiosis, we used the pool of cassava genes that were significantly differentially expressed between the two AMF treatments and the non-mycorrhizal treatment to build cassava gene modules. On the other hand, because AMF can only be grown in presence of the host plant, we could not select AMF genes that were only involved in the symbiosis. We, therefore, selected all the *R*. *irregularis* genes in the dataset in order to build *R*. *irregularis* gene modules. We used the default WGCNA ‘step-by-step network construction’ analysis to build modules. We first calculated the adjacency between genes and constructed a topological overlap matrix. We then produced a hierarchical clustering tree with the dissimilarity of the topological overlap matrix and we selected the modules by using the dynamic three cut standard method. Finally, we merged similar modules by calculating the module eigengenes, clustering them and assigning a distance threshold. The parameters used were soft-power of 12 and minimum module size of 50 genes. We merged the modules with a distance threshold cut of 0.1.

We associated the modules between the two organisms by measuring the Pearson correlation between cassava module eigengenes and *R*. *irregularis* module eigengenes using the default WGCNA ‘relating modules to external information’ analysis. We calculated the gene significance (Pearson correlation) for each gene to the corresponding correlated module (e.g. gene1-cassava vs. module1- *R*. *irregularis* etc.) and we calculated the connectivity (membership) of each gene to its module. We used this information in order to obtain a set of ‘key genes’ of a module. These are defined as the genes that are the most correlated to modules of the other organism (in the top 10% quantile) and that also displayed a high module membership (>0.8). All the module pairwise comparisons result in a dataset of ‘key genes’ that are highly connected within each module and highly correlated to modules of the partner organism.

We also performed the same analyses as before (‘relating modules to external information’) to correlate the cassava and *R*. *irregularis* modules to fungal and plant growth variables. We used the same parameters and threshold cut-off for these comparisons.

## Results

### Plant growth and fungal colonization

We observed almost zero colonization levels of the cassava plants in the mock-inoculated treatments, confirming that these treatments were not colonized by AMF (Supplementary figure 2a). The cassava cultivars displayed significantly different levels of AMF colonization, but colonization by isolates B1 and DAOM17978 was not statistically different (Supplementary Figure 2a; Supplementary Table 2a). We observed an overall effect of the plant cultivar on all the quantitative trait measurements, but we did not observe an overall effect of the fungal identity on the quantitative trait measurements (Supplementary Table 2). Despite this, we observed that the dry weight of cultivar COL2215 was significantly higher when inoculated with isolate B1 compared to the mock control (Supplementary Figure 2b; Supplementary Table 2b).

**Figure 2.**
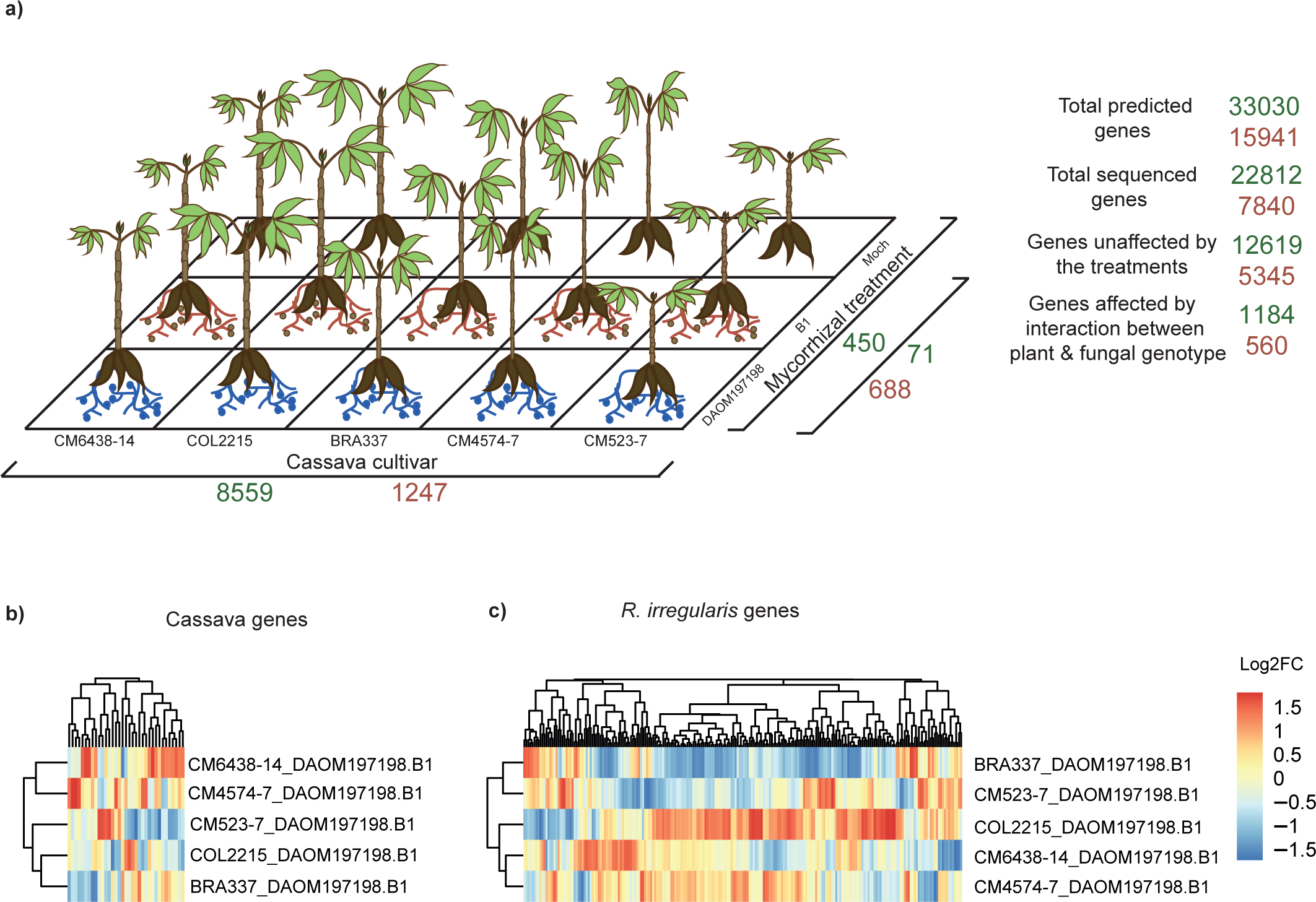
Variation in cassava and *R*. *irregularis* gene transcription. **(a)** Number of gene transcripts in the different treatments. Cassava genes are reported in green. *R*. *irregularis* genes are reported in brown. **(b)** Cassava genes with a log2 fold change higher than 2 between plants inoculated with isolates DAOM17978 or B1. **(c)** *R*. *irregularis* genes with a log2 fold change higher to 4 between isolates DAOM17978 or B1 living with different cassava cultivars. Genes (columns) and fold change by cassava cultivar (rows) are clustered by similarity. Gene names can be found in Supplementary table 4d and Supplementary table 5d.

### Activation of plant ‘mycorrhizal toolkit’ genes in presence of AMF

In order to test the robustness of the plant RNA-seq data, we identified 12 cassava orthologs of genes that are known to be differentially transcribed in response to AMF inoculation in *Medicago trunculata* (Supplementary note S6). As expected, this set of genes, which is part of the conserved plant “mycorrhizal toolkit”, was clearly differentially transcribed between the mycorrhizal and non-mycorrhizal treatments in all the cultivars (Figure 1; Supplementary Table 3).

### Cassava gene transcriptional response to the identity of the fungal partner is plant genotype specific

A large number of cassava gene transcripts were recovered from the roots, representing 69% of the total predicted gene number (Figure 2a). Of the total number of transcribed genes, 10193 genes (44.6%) were significantly differentially affected by plant genetic variability or the mycorrhizal status of the plants, or both. We did not observe any sequencing bias among the samples, as the number of different cassava gene transcripts did not differ significantly among mycorrhizal treatments, or among cassava cultivars (Supplementary Figure 3a-c), indicating that the dataset was reliable for detecting true transcriptional differences. The transcription of 8559 cassava genes differed significantly among the genetically different cassava cultivars but was not affected by the mycorrhizal symbiosis (Figure 2a; Supplementary Table 4a). Only a small number of cassava genes (450 genes) were influenced by the mycorrhizal treatments in a conserved way, meaning that they were not differentially transcribed among cassava cultivars; with most differences occurring between the mock-inoculated and mycorrhizal plants (Figure 2a; Supplementary Table 4b). Only 71 cassava genes were differentially transcribed in plants inoculated with the two different fungal isolates and that also showed consistent transcription patterns among all cassava cultivars (Figure 2a; Supplementary Table 4b). In contrast we observed a strong genotype x genotype interaction on gene transcription as the transcription of a surprisingly large number of cassava genes (1184 genes) was significantly affected by the combination of the genetic identity of both the plant and the mycorrhizal fungus (Figure 2a; Supplementary Table 4c). This means that transcription of those cassava genes were influenced by the presence of the fungus but that the magnitude of this response to the fungus differed quantitatively and in direction among the plant cultivars. From this number, thirty-five cassava genes, displaying large fold changes in transcription between the two fungal treatments, DAOM 197198 and B1, responded in a significantly different way among the cassava cultivars (Figure 2b; Supplementary Table 4d). Many genes involved in the response to biotic and abiotic stresses, categorized as being involved in signalling, secondary metabolites, proteolysis and cell wall constituents were differentially transcribed in a different direction among cultivars inoculated with the different fungal isolates (Supplementary Figure 4a).

**Figure 3.**
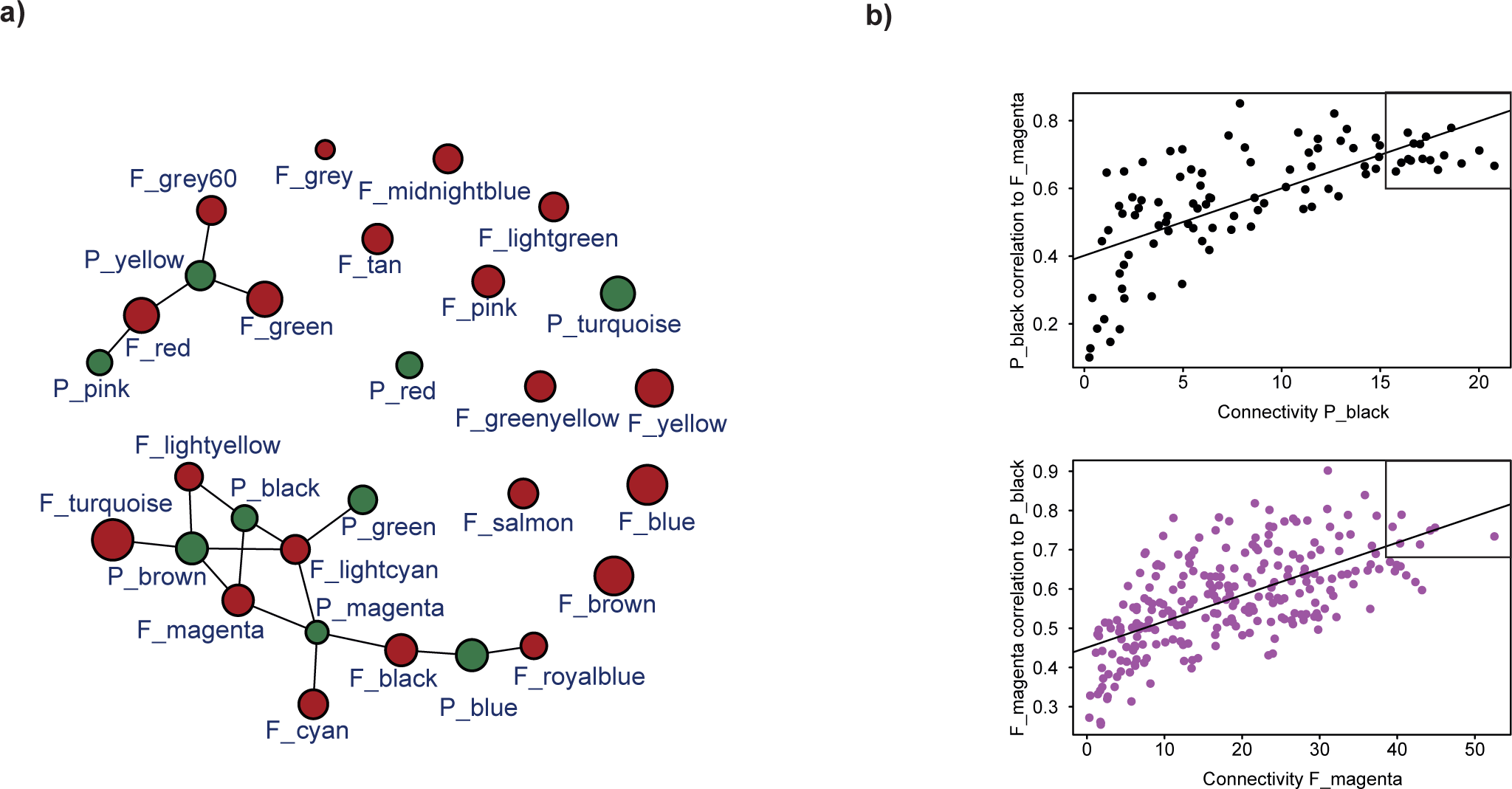
Gene co-expression network analysis. **(a)** Network of *R*. *irregularis* and cassava modules. Highly correlated modules (*p-value* <0.001) are linked with a line. Brown and green circles represent *R*. *irregularis* and plant modules, respectively. **(b)** Example of the identification of key genes between two correlated modules for one plant gene module (module P black) and one fungal gene module (module F magenta). Each dot represents one gene and ‘key genes’ (the top right of each graph) are the most correlated to modules of the other organism (top 10% quantile) and that also displayed a high module membership (>0.8).

**Figure 4.**
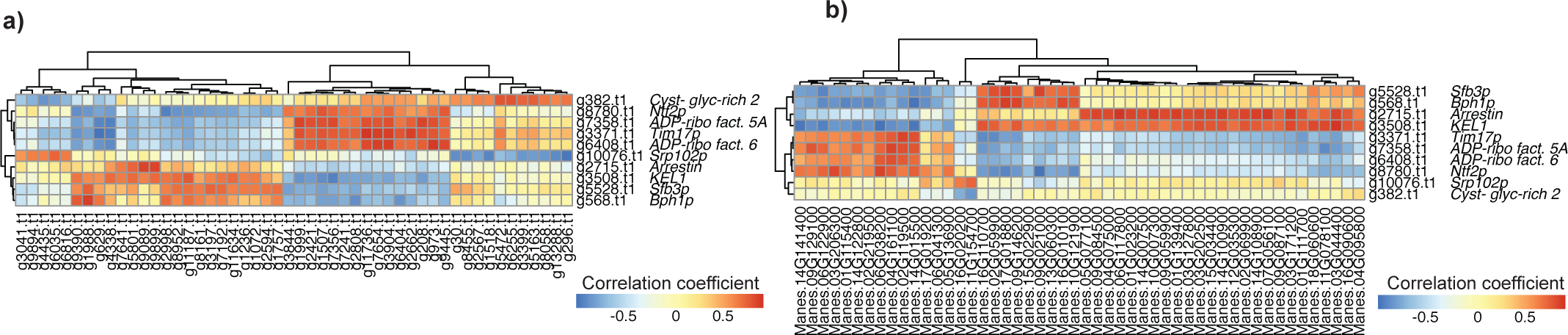
Correlogram of *R*. *irregularis* genes involved in the secretory pathway. Secretory pathway genes correlated with: **(a)** 50 fungal genes randomly sampled from 317 *R*. *irregularis* key genes. **(b)** 50 cassava genes randomly sampled from 120 cassava key genes. Each square represents the Pearson correlation coefficient between the genes. Genes in rows and columns were clustered by the similarity of their correlation coefficient. We grouped all the ‘key genes’ by functional categories and correlated them to all the ‘key genes’. All the correlograms made in this study can be found in Supplementary Table 7.

### Fungal gene transcription is strongly affected by genotypic variation in cassava

The 7840 recovered fungal gene transcripts represented 49% of the predicted gene number (Figure 2a). Very few fungal transcripts were recovered from samples that were mock-inoculated and read number did not influence the number of gene transcripts, showing that there was no bias in the fungal transcriptome data (Supplementary Figure 3a-c). Genetic variation among the cassava cultivars had a strong effect on fungal gene transcription, where 1247 fungal genes were differentially transcribed among cassava cultivars (representing 69% of the differentially transcribed fungal genes; Figure 2a; Supplementary Table 5a). Differential transcription of a smaller number of genes (688 genes) occurred between the two fungal isolates irrespective of the cassava cultivar (Figure 2a; Supplementary Table 5b). We also observed a genotype x genotype interaction as a substantial number of fungal genes (560 genes, representing 31% of the differentially transcribed fungal genes) were differentially transcribed between fungal isolates but in a different direction or magnitude in different cassava cultivars (Figure 2a; Supplementary Table 5c), showing lack of conservation in their expression. Of these 560 genes, a high number of fungal genes (269 genes) showed large fold changes in transcription between the two fungi but differed strongly in transcription among cassava cultivars (Figure 2c; Supplementary Table 5d). Genes involved in the auxin pathway, signalling, glutathione-S-transferase, cell wall constituents, proteolysis and MAP kinases and many genes involved in secondary metabolite production were significantly transcribed in different directions in response to cassava genetic variability (Supplementary Figure 4b).

### Gene co-transcribed networks as a tool to decipher molecular interactions of the mycorrhizal symbiosis

The measurement of plant and fungal gene transcription, and its variability within the same experiment caused by different treatments, allowed us to evaluate co-ordinated molecular interactions between cassava and the fungi by identifying genes of the plant and the fungus that were highly correlated. We used the variance in gene transcription that resulted from the experimental design in order to identify co-transcribed plant and fungal gene clusters. The weighted gene co-expression network approach (WGCNA) allowed us to identify 9 and 20 modules of co-transcribed plant and fungal genes, respectively (Figure 3a, Supplementary Figure 5ab; Supplementary Table 6ab). Within the identified modules, we detected 7 cassava gene modules and 10 fungal gene modules that were correlated to at least one module of the partner species (Figure 3a; Supplementary Figure 5c). To confirm that the modules were biologically meaningful we randomly attributed genes to cassava and fungal modules and attempted to construct a new gene co-transcribed network. None of these modules corresponded to significant gene ontology terms and yielded no network (Supplementary Figure 6). Thus, the observed gene co-transcribed network was unlikely to have occurred by chance. We calculated the gene connectivity within the modules in all genes found in modules that were correlated to a partner module and the gene’s correlation to the partner module. This allowed us to detect ‘key genes’ within a given module (for an example of correlation between 2 modules see Figure 3b). Within all the plant and fungal correlated modules, we found 120 cassava and 317 *R*. *irregularis* ‘key genes’ in total that were highly representative of the modules and highly correlated to a counterpart partner module (Supplementary Table 6cd).

**Figure 5.**
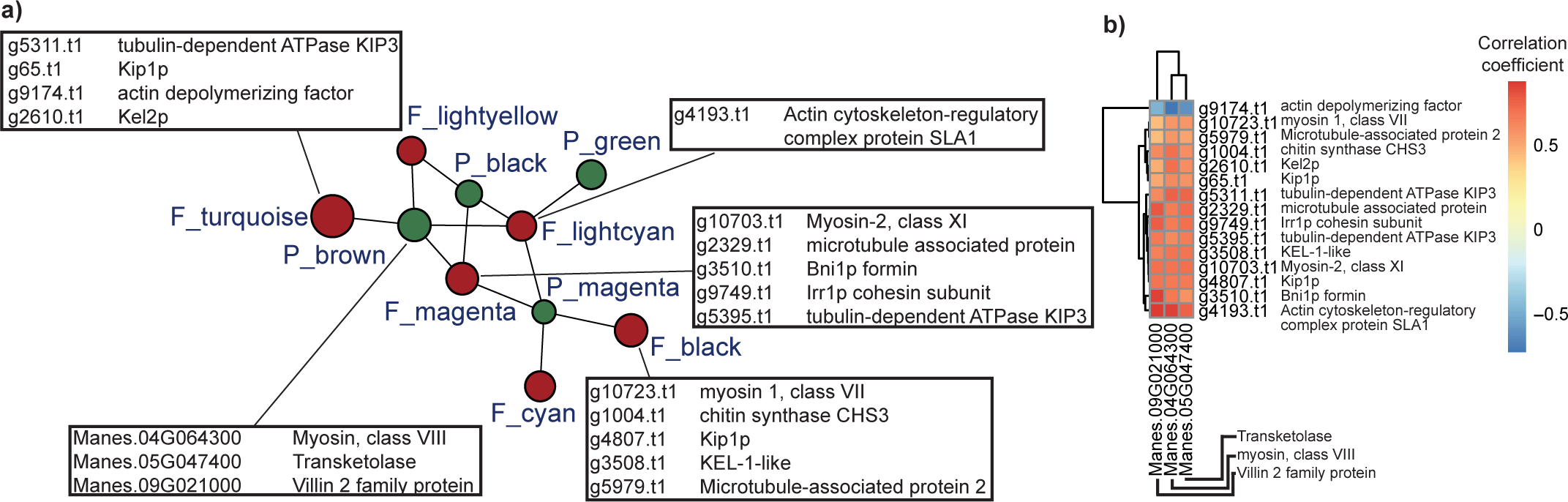
Cellular organisation gene transcription in the AM symbiosis. **(a)** *R*. *irregularis* and cassava genes involved in cellular organisation. We show genes involved in cellular organisation that were highly correlated with gene transcription in the partner organism. **(b)** Correlogram of the *R*. *irregularis* and cassava genes involved in cellular organisation. The information about gene name and function of the ‘key genes’ is found in Supplementary Table 6.

**Figure 6.**
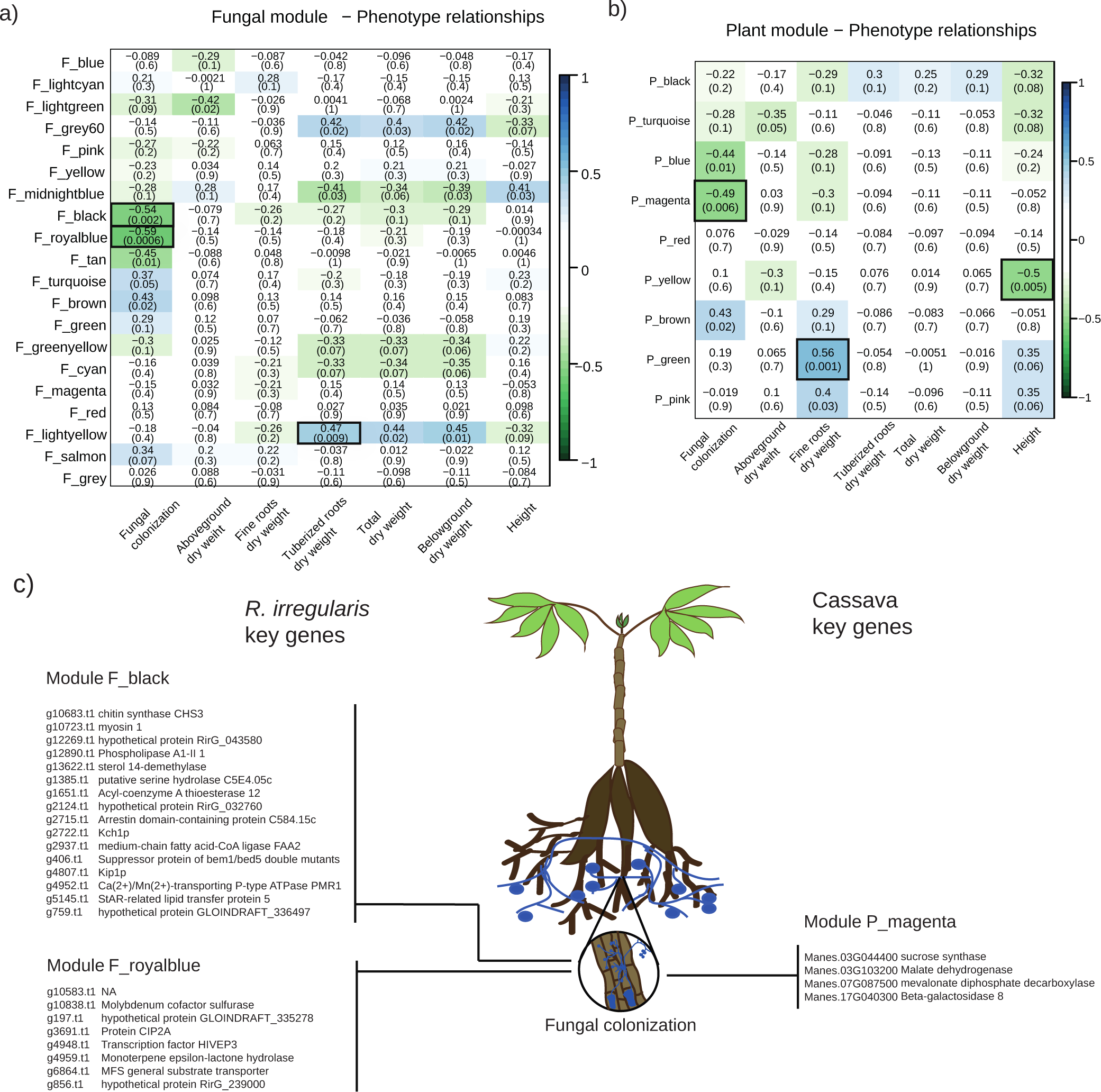
*R. irregularis* and cassava modules correlated with quantitative phenotypic traits. Correlation matrix between **(a)** the cassava modules, or **(b)** the *R*. *irregularis* modules (rows) and the quantitative traits (columns). We report the Pearson coefficient and its *p*-value. Highly positive correlations are shown in dark blue and highly negative correlations are shown in dark green. **(c)** Overview of modules and the ‘key genes’ highly correlated to the quantitative trait fungal colonization. (Gene function and correlation score can be found in Supplementary table 6; for multiple comparison correction threshold refer to Supplementary note S9).

This approach allowed us to detect several genes with different functions that were correlated with genes within the same organism and the partner organism. For example, we found 10 key genes involved in the fungal secretory pathway that displayed either highly positive or highly negative correlations to other fungal and plant genes. This suggests that the function of this group of genes is heterogeneous, and reveals that approximately half of the genes involved in the secretory pathway are antagonistic (Figure 4; Supplementary Table 6cd). Notably, 8 of these genes were either strongly positively or negatively correlated with genes that are expected to play an important role in the symbiosis such as plant phosphate transporters and sucrose synthases, suggesting an indirect correlation of the secretory pathway in the genetic reprogramming of the symbiosis (Supplementary Table 7). We also found that several key fungal and plant genes in the network represented genes involved in the cytoskeleton and cellular organization. These genes were highly positively correlated between the two organisms (Figure 5; Supplementary Table 6d).

### Transcription of plant and fungal genes correlated with fungal colonization

We applied the WGCNA method to detect plant and fungal genes that were correlated with quantitative traits of either fungal or plant growth. We found 3 plant and 3 fungal gene modules that were correlated with quantitative traits. Within these modules, we found that two *R*. *irregularis* modules and one cassava module correlated with fungal colonization (Figure 6). The fungal module “F_black” and “P_magenta”, where correlated with fungal colonization, as well as being highly co-correlated each other (Supplementary Table 6d). Notably, fungal chitin synthase was highly correlated with plant sucrose synthase transcription. In addition, both of these genes were highly correlated with fungal colonization (Figure 6).

## Discussion

In this study, we evaluated plant and fungal quantitative growth responses and transcriptional patterns of both partners in response to intraspecific genetic variation. The main finding of this study is that contrary to what is already known about the highly conserved plant molecular responses allowing colonization of roots by mycorrhizal fungi, we found a very large plant genotype x fungal genotype interaction on gene transcription in both organisms, involving an unexpectedly large number of plant and fungal genes. What this means is that genetic reprogramming occurred in cassava roots in response to AMF, but that the magnitude and direction of the transcriptional reprogramming was not conserved, differing greatly among cassava cultivars and fungal genotypes (genotype x genotype interaction). In addition, the genetic variation, used as treatments in this experiment, resulted in considerable quantitative differences in gene transcription levels in the partner organisms that allowed us to detect genes that were highly correlated between the two organisms, as well that were correlated to quantitative growth traits. Such detection of plant-AM fungal gene networks has not been possible in previous studies using simpler experimental designs. The ecological implications of these findings are discussed below.

### Conserved and non-conserved molecular reprograming in the mycorrhizal symbiosis and its ecological relevance

One of the remarkable features of the last two decades of investigation into plant genes involved in the arbuscular mycorrhizal symbiosis is the fact that a very large number of the plant genes that are involved in the establishment of arbuscular mycorrhizal fungi inside the roots are highly conserved across a large part of the plant kingdom, including basal plant groups (Delaux Ané et al). Coupled with approaches using plant mutants, where disruption of these genes leads to impaired development of the fungus inside the roots, this has lead to the assumption that the molecular basis for the interaction between plants and AM fungi is highly controlled by the plant and also expected to be highly conserved among plants that are capable of forming the mycorrhizal symbiosis. The results of our study clearly showed that a large number of these plant genes were upregulated across all treatments where the fungus was present and irrespective of plant or fungal genetic variation, supporting the hypothesis that fundamental genes involved in the physical establishment of the fungus inside the host root are indeed conserved.

Despite this conservation of genes involved in the establishment of the symbiosis, our study shows that following symbiosis establishment, the molecular reprograming that occurs in both the plant and the fungus is highly plant and fungal genotype specific and differentially affects the transcription of a very large number of genes in both organisms. In fact, 72% of the cassava genes that were affected by the mycorrhizal symbiosis were transcribed in a different direction or magnitude among the cassava cultivars and, thus, responded in a non-conserved manner. While previous molecular studies of the symbiosis have shown conservation across plant species, families and orders, we show that the molecular reprogramming during symbiosis is very highly affected by genetic variability, even among closely-related genotypes of one plant species and one fungal species, indicating the high degree of molecular complexity of the interaction following symbiosis establishment and a lack of conservation.

In soil, a mycorrhizal fungus repeatedly encounters roots of different plants (different plant species and different genotypes within plant species). *Vice versa*, in soil, a plant encounters different mycorrhizal fungal species and genotypes of a species. Ecological studies show that the outcome of the different combinations of plant and fungal genotypes can lead to very variable outcomes in terms of growth benefits for both individuals and the outcome (at least in terms of growth) is not always positive. In this study, plant and fungal growth varied as a result of the genetically variable treatments and yet upregulation of the conserved repertoire of genes that allowed the establishment of the symbiosis occurred irrespective of whether the establishment was beneficial or not. Based on previously published ecological studies, coupled with the results of this study, we propose that underlying the highly conserved molecular regulation allowing symbiosis establishment, a highly non-conserved genetic reprograming occurs in the plant and the fungus that is highly dependent on the genotype of the partner, even within species. The implications of this are that in order to understand the molecular interactions between the plant and the fungus that determines the fitness benefits afforded by associating with a given partner requires in depth studies that consider genetic variability of both plants and the fungus, even at the level of intraspecific diversity.

In this study we observed a non-conserved response of the cassava cultivars to the AMF isolates, particularly in genes involved in plant defence, signalling, cell wall components and proteolysis. This implies that the host plant could elicit a different strategy to the colonization by different AMF genotypes and we hypothesize that depending on the evolutionary history and adaptation of different genotypes, their capacity to perceive and respond to the fungus differs.

### Conserved and non-conserved fungal transcriptional responses to cassava and its genetic variation

We were able to separate AM fungal gene transcription patterns that were conserved across plant genotypes and those that were non-conserved; varying in direction and magnitude of transcriptional patterns as a result of plant genetic variation. Because almost all published transcriptome studies on the AM symbiosis use an experimental design with one inoculated plant genotype compared to a mock inoculated control, this does not allow the detection of differential AMF gene transcription as the control treatment cannot contain fungal transcripts. Thus, this is the first investigation revealing the conserved *versus* non-conserved genetic reprograming in the fungus during the mycorrhizal symbiosis. The non-conserved fungal genes, among others, were related to the auxin hormone pathway, which is known to be involved in the development of arbuscules [36], glutathione-S-transferases that could be involved in heavy metal binding [37] and MAP Kinases which are known to play a role in the response to biotic and abiotic stresses [38, 39]. These results demonstrate that the two AMF strains could have different symbiotic strategies and that these strategies could be modified in presence of a different host plant. Evidence of genotype x genotype interactions on fungal quantitative traits has been already described in AMF [40, 41]. In this study we confirmed, by measuring gene transcription, that genotype x genotype interactions are a common feature that should be addressed to better understand the molecular interactions between both partners.

In order to understand the molecular mechanisms of beneficial AMF-plant interactions, a more extensive comparison needs to be made between plants exhibiting benefit from AMF colonization with plants that do not respond, and using different plant models. The identification of a ‘beneficial plant mycorrhizal toolkit’ would be of great agronomic and ecological interest and would represent a milestone in the applied AMF research.

### Gene co-expression networks as a tool to decipher molecular interactions in the mycorrhizal symbiosis

The dual RNA-seq approach on AMF colonized plant roots allowed us to identify fungal and plant transcripts that were co-expressed. Dual RNA-seq is an emerging approach that has allowed researchers to decipher interactions between hosts and pathogens [42-45]. The WGCNA approach decreases the complexity of such data sets by clustering gene transcripts and allowing the identification of co-expression patterns between the two organisms, as well as to quantitative traits of the organisms [46]. In this study, we only focussed on the network existing between plant and AM fungal genes, which allowed us to get biological insights into the crosstalk in the mycorrhizal symbiosis. Our study showed that an extensive network of co-expressed gene exists between plant and AM fungal genes. We identified several ‘key’ plant and fungal genes related to cytoskeleton formation and cell organization that were highly co-expressed within and between both organisms, with a strong positive co-expression of between actin and myosin genes, and a strong negative correlation between actin genes and an actin depolymerisation factor, indicating that plausible biological processes could be detected with this network approach.

While the process of arbuscule formation inside the plant cells has been widely described [47], there is little data focusing on the cellular organisation and cytoskeleton changes occurring at later stages of the symbiosis. Cell wall organisation has been studied histologically showing that plant actin filaments and microtubules appear around arbuscular branches, but disappear at arbuscule senescence [48]. In addition, plant microtubules in cells adjacent to fungal structures are also modified, indicating signal exchange between the symbionts prior to fungal penetration [49]. However, there is little information about the changes in cellular organisation in the fungal structures, possibly, because more emphasis has been placed on plant structures [50]. Our study shows that there is a strong correlation between genes involved in the cell organisation of the plant and the fungus, which suggest that these processes are probably co-orchestrated.

### Correlation of gene-transcription with fungal colonization

We found a strong correlation between fungal colonization of the roots and one plant gene module and two fungal gene modules. Furthermore, both plant and fungal gene modules were highly correlated with each other. Within these modules, notably, we observed a fungal chitin synthase and a plant sucrose synthase that correlated to the fungal colonization. Plant sugar transport, is important for allocation carbohydrates to the fungal symbiont [51]. The co-expressed plant sucrose synthase observed in this study has a conserved domain (TIGR02470), the role of which is to make sucrose available at the cell wall. We hypothesize that the allocation of carbohydrates from the plant to the fungus also has an effect on the fungal colonization phenotype observed in this experiment. We also observed a correlation of the fungal chitin synthase with fungal colonization. Chitin is used by plants as a signal that triggers plant defence [52]. The correlation between the chitin synthase, sucrose synthase and the fungal colonization, suggests that chitin synthase could also have a role in the maintenance of the fungi inside the plant root. We found other ‘key genes’ that correlated with fungal colonization but their function is less well described. In addition, we classified by functional categories, the ‘key genes’ found in each module, and we were able to identify strong correlations among and between plant and fungal genes in functional categories (Supplementary table 7). The study in detail of the gene-gene interactions on all the other functional categories is out of the scope of this study. However, the list of genes highlighted in this study could be a starting point to better understand the molecular basis of the interaction between the host plant and the AMF symbiont.

### Experimental design in elucidating ecologically relevant molecular interactions between plants and AMF

Our study is unique because the experimental treatments of varying plant and fungal genotype allowed us to identify molecular interactions between the plant and the fungus that have not previously been achieved with a conventional mycorrhizal plant *versus* non-mycorrhizal plant comparison. Firstly, this allowed us to investigate the role of genetic variation in the molecular interplay between plants and AM fungi and quantify the conserved *versus* non-conserved component of these molecular interactions; Second, it allowed us to study variation in fungal transcripts which is completely novel; Third, because genetic variation was used as replicated experimental treatments, the resulting variation in plant and fungal gene transcription allowed us to identify a robust co-expression network that exists between the plant and the fungus. None of these results would be identifiable or quantifiable without more complex designs than a simple mycorrhizal versus non-mycorrhizal comparison. While we are fully aware of the enormous advances that have been made in the role of plant genes involved in the establishment of the symbiosis, by the use of plant mutants with impaired fungal development, and that our study of gene transcription cannot experimentally reveal the role of individual genes, our study strongly demonstrates the quantitatively large genotype intraspecific variation in the molecular interactions in the mycorrhizal symbiosis. In order to fully understand the molecular interactions between plants and mycorrhizal fungi that are ecologically relevant and leading to change in plant or fungal fitness, researchers will have to embrace genetic variation in the partners to fully understand these interactions and not rely on simple experimental designs.

## Acknowledgements

We thank Jeremy Bonvin for help culturing *R*. *irregularis* and cassava *in-vitro* plants, Tania Wyss for help in fungal culturing and plant harvesting and Nicolas Ruch for help in the greenhouse. We thank the International Center for Tropical Agriculture (CIAT) for providing cassava cultivars. All bioinformatics computations were performed at the Vital-IT (http://www.vital-it.ch) Center for High Performance Computing of the Swiss Institute of Bioinformatics. All sequencing was performed at the Genomic Technologies Facility (GTF) of the University of Lausanne. This study was funded by the Swiss National Science Foundation (Grant number: 31003A_162549 to IRS). Sequence reads were deposited in the NCBI SRA database (BioProject Accession Number: PRJNA400637).

